# Production of recombinant heterotrimeric mini-procollagen I and homotrimeric mini-procollagen II reveals new cleavage sites for BMP-1

**DOI:** 10.1101/2022.11.10.516045

**Authors:** Natacha Mariano, Cindy Dieryckx, Agnès Tessier, Jean-Baptiste Vincourt, Sandrine Vadon-Le Goff, Catherine Moali

## Abstract

The proteolytic conversion of soluble procollagens into mature collagen monomers is a critical step to decrease their solubility and trigger collagen fibril formation. In the case of collagens I, II and III, this maturation process is driven by several extracellular metalloproteinases such as BMP-1, tolloid-like proteinases, meprin α, meprin β, ADAMTS-2 and ADAMTS-14 but the extensive characterization of these proteolytic events has been hampered by the lack of recombinant procollagens. We previously reported the production and partial characterization of recombinant homotrimeric proteins derived from procollagen III (mini-procollagens III) and, in this study, we describe how we have extended this previous work to the production of heterotrimeric mini-procollagen I and homotrimeric mini-procollagen II. These mini-procollagens include truncated triple helices and intact C-telopeptide and C-propeptide domains and were produced in suspension in HEK293-F cells with yields ranging from 2.5 mg/L to 10 mg/L after purification. They proved very useful tools to analyze the effect of calcium on the stability of the procollagen C-terminal region and to compare the procollagen C-proteinase activity of BMP-1 on the three major fibrillar procollagens or their ability to interact with various partners such as PCPE-1. Using mass spectrometry to map BMP-1 cleavage sites on the mini-procollagens, we confirmed all previously described sites but also revealed two additional cleavage sites in the α1 chain of procollagens I and II. This result shows that the mini-procollagen toolkit offers a broad range of perspectives to make functional studies but also possibly structural analyses or to develop drug screening assays.

## Introduction

Procollagens are the precursor forms of mature fibrillar collagens. Both precursor and mature collagens share a central triple helical domain, made of more than 300 Gly-Xaa-Yaa repeats flanked by N-and C-telopeptide regions, but procollagens also contain additional domains at both extremities, called N-and C-propeptides (1, 2). These domains maintain procollagens in the soluble form until they undergo proteolytic maturation by procollagen N-and C-proteinases (PNPs and PCPs), a critical step to trigger fibril formation. Major fibrillar collagens are types I, II and III which are by far the most abundant protein components in connective tissues such as skin dermis (3), corneal stroma (4), cartilage (5), tendons (6) or bones (7). While procollagen I is predominantly found in tissues as a heterotrimer consisting of two α1 chains and one α2 chain, procollagens II and III are homotrimers composed of three identical α1 chains.

A better understanding of the proteolytic conversion of procollagens I, II and III, taken individually or as defined mixtures, into mature collagens would give important insights into how tissue-specific properties of collagen fibrils are established and maintained (2). It would also help to design appropriate therapeutic strategies to control defective or excessive collagen deposition, for example in impaired wound healing, fibrosis or in genetic diseases. So far, this type of functional and structural studies has been hampered by the difficulty to obtain large amounts of the various procollagens.

Native procollagen I can be successfully enriched in milligram amounts from the supernatant of skin or tendon fibroblasts cultivated in the presence of ascorbic acid (8, 9) but partial proteolysis by endogenous PNPs and PCPs often occurs, making separation of processed and unprocessed forms challenging. The same applies to the extraction of native procollagen III from rat skin (10). Recombinant expression of full-length procollagens I-III has also been achieved with very high yields in yeast *Pichia pastoris* through the co-expression of recombinant prolyl-4-hydroxylase to ensure proper proline hydroxylation (11, 12). However, in this system, procollagens are not secreted and accumulate in the endoplasmic reticulum and the demonstration that their successful extraction is possible remains to be done. Only the extraction of pepsin-released triple helical domains has been reported until now. More common production systems for procollagens are now mammalian cells such as HT1080 or HEK293 which have endogenous prolyl-4-hydroxylase activity (13). We have used HEK293-EBNA cells in the past for procollagen III expression (14) but we observed substantial protein loss during purification and recurrent contamination by low-molecular weight proteins which proved to be histones.

Based on the numerous pitfalls of the production and purification of full-length procollagens, we and others have developed recombinant mini-procollagens which correspond to truncated versions of human procollagens and can be tailored for various applications. Especially, we previously designed and characterized several mini-procollagens derived from collagen III which, in addition to the C-propeptide that drives the trimerization process, included N-terminal extensions of variable lengths. These constructs proved extremely useful for the characterization of metalloproteinases with procollagen C-proteinase activity, namely BMP-1 (bone morphogenetic protein-1)/tolloid-like proteinases (i.e. BMP-1, mTLD, mTLL-1 and -2) (14, 15), meprins α and β (16), ADAMTS (A Disintegrin And Metalloprotease with Thrombospondin motifs) -2 and -14 (17), but also of the regulation of these maturation events by PCPE-1 (procollagen C-proteinase enhancer protein-1). For example, successive truncations of a mini-procollagen III (Mini-III), initially composed of the 33 most C-terminal GXY triplets, the native C-telopeptide and the native C-propeptide found in human procollagen III, down to a construct containing only 2 residues and a tag in addition to the C-propeptide allowed us to prove that BMP-1/tolloid-like proteinases (BTPs) can still cleave this short protein and that PCPE-1 enhances procollagen processing by BTPs through C-propeptide binding (14, 15). The shortest forms also proved very useful for structural analysis by SAXS and X-ray crystallography (18–20). All the described proteins were produced either through stable transfection in HEK293-EBNA or through transient transfection in HEK293-T or -F cells.

In the present manuscript, we describe how we have built on this previous experience to address the challenge of producing a heterotrimeric human mini-procollagen I, something which has never been done before. We already knew that it was possible to generate recombinant heterotrimeric C-propeptide I in HEK293 cells and to separate it from (α1)_3_ homotrimers through the introduction of the tag on the α2 chain (21, 22) and we decided to use a similar strategy to extend the length of the protein in the N-terminal direction. In parallel, we also produced a mini-procollagen II to complete the mini-procollagen toolkit. The production of these proteins and their preliminary structural and functional characterization is described below.

## Results

### Production and purification of mini-procollagens

The new mini-procollagen I (Mini-I) and mini-procollagen II (Mini-II) constructs (Fig. 1A) encompassed the last GXY triplets (between 11 and 33) found in native human sequences, stabilized by additional GPP repeats at the N-terminus, and full-length C-telopeptide/C-propeptide sequences (Fig. S1). A 6His tag was added at the N-terminus of Mini-II chain to facilitate purification and a Twinstrep tag was specifically incorporated in the α2(I) chain of Mini-I to allow the separation of [(α1)_2_]α2 heterotrimers from (α1)_3_ homotrimers (21, 22). As previous work has showed that N-glycosylation of C-propeptides is not required for protein folding in normal conditions (18, 22, 23), we also mutated the N-glycosylation sites (N1365 in the α1(I) chain and N1267 in the α2(I) chain) into alanine in Mini-I.

**Figure 1:**
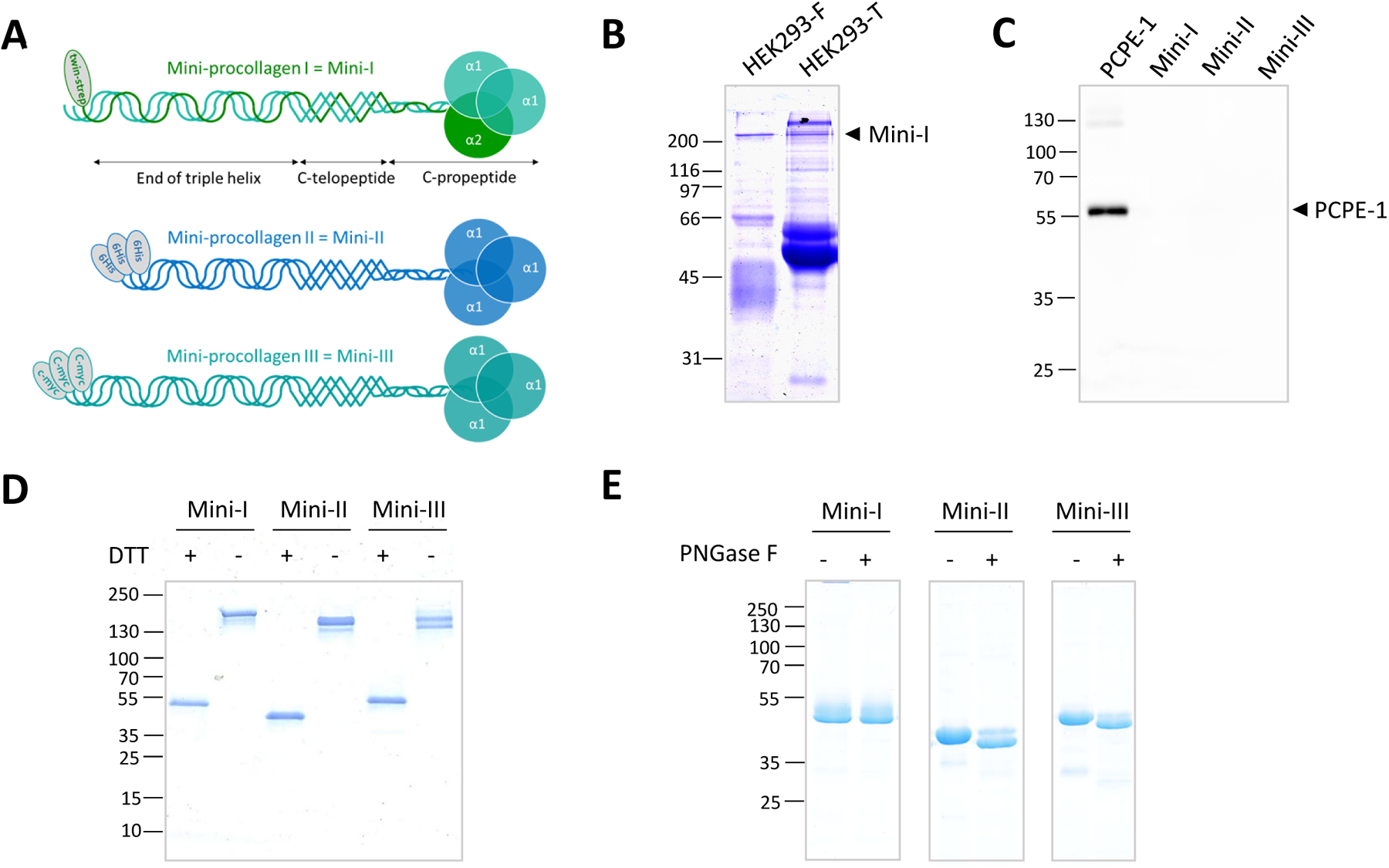
Design and production of mini-procollagens. **(A)** Mini-procollagens are composed of native C-propeptides, native C-telopeptides, truncated triple helices and N-terminal tags. **(B)** SDS-PAGE analysis of the conditioned medium of HEK293-T and -F after transfection with pHLsec plasmids encoding the α1 and α2 chains of Mini-I (10 % acrylamide gel, non-reducing conditions, Coomassie Blue staining). **(C)** Immunoblot showing that PCPE-1 is not present in final preparations of Mini-I, -II and -III. One microgram of each mini-procollagen was loaded on a 10 % acrylamide gel (non-reducing conditions) and recombinant human PCPE-1 (20 ng) was used as positive control. **(D)** Trimer formation is evidenced by a shift in molecular weight upon DTT reduction (4-20 % acrylamide gradient gel, InstantBlue staining). **(E)** Removal of N-glycosylations in Mini-II and -III by PNGase F leads to a shift in molecular weight that is not observed in Mini-I, showing the successful ablation of the N-glycosylation sites in the latter protein (4-20 % acrylamide gradient gel, reducing conditions, InstantBlue staining). The gel is representative of two independent experiments.

For protein production, we tested two validated culture systems allowing the transient transfection of either adherent cells (HEK293-T cells) or cells grown in suspension (HEK293-F cells). Although more expensive to maintain due to the use of proprietary defined media, the HEK293-F cells offer the advantage to grow in the absence of serum whereas HEK293-T cells are usually transfected in the presence of 2 % serum, leading to more contaminants and to a lower reproducibility. Production yields in the two systems appeared to be similar (Fig. 1B) and we chose to work with the HEK293-F cells and pHLsec plasmids (24) for transient transfection.

Purification was achieved in two steps, affinity purification through the tag and size exclusion chromatography (SEC), as described in Materials and Methods. Typical yields after purification were 2.5 mg/L of culture medium for Mini-I and 10 mg/L for Mini-II, to be compared with 1 mg/mL for Mini-III produced in HEK293-EBNA cells. Importantly, Mini-III purification required one preliminary step (Affiblue gel or heparin sepharose) to remove major serum contaminants and one additional passage through anti-PCPE-1 sepharose to get rid of the residual PCPE-1 that is secreted in relatively high amounts by HEK293-EBNA cells and remains bound to Mini-III. In contrast, HEK293-T and -F cells were found to express less PCPE-1 and no endogenous PCPE-1 was found to co-purify with Mini-I and Mini-II (Fig. 1C).

Proper trimerization of mini-procollagens was demonstrated by SDS-PAGE, in reducing and non-reducing conditions (Fig. 1D). The observed bands were in good agreement with the expected sizes of the respective monomers and trimers. However, the heterotrimeric nature of Mini-I could not be confirmed at this stage because both α1 and α2 chains co-migrated due to the integration of the relatively long Twinstrep tag in the α2 chain that is normally shorter than the α1 chain. Additional information on this aspect will be described below. Finally, absence of N-glycosylation sites in Mini-I was confirmed by the absence of shift after incubation with PNGase F while Mini-II and -III displayed a lower apparent molecular weight after the same treatment (Fig. 1E), in agreement with the retention of their N-glycosylation site.

### Stability of mini-procollagens in the absence and presence of calcium

Calcium is known to play a major role to stabilize C-propeptide structures (18, 21, 22) and we next analyzed the impact of calcium removal on mini-procollagen stability using Nano-DSF (Differential Scanning Fluorometry) (Fig. 2). Denaturation curves were recorded for each mini-procollagen, either in HEPES buffer containing 5 mM calcium chloride or after addition of 20 mM EDTA. First and importantly, there was no sign of aggregation for any of the mini-procollagens in either condition. Then, we observed that, in the presence of calcium, Mini-I and Mini-II showed two clearly defined transitions corresponding to distinct unfolding events. Mini-II also seemed significantly more stable than Mini-I with a melting temperature (Tm) of the first denaturation event that was more than 10 °C higher. For Mini-III, only one peak was clearly visible, with a Tm of 56.3 °C, in the range of the lowest transition of Mini-I and -II, but a second event also seemed to appear above 90 °C. This suggested a higher thermal stability for the specific domain of Mini-III affected by the second transition, that could be due to the presence of an additional inter-chain disulfide bond at the C-terminal extremity of the triple helix (absent in the other mini-procollagens).

**Figure 2:**
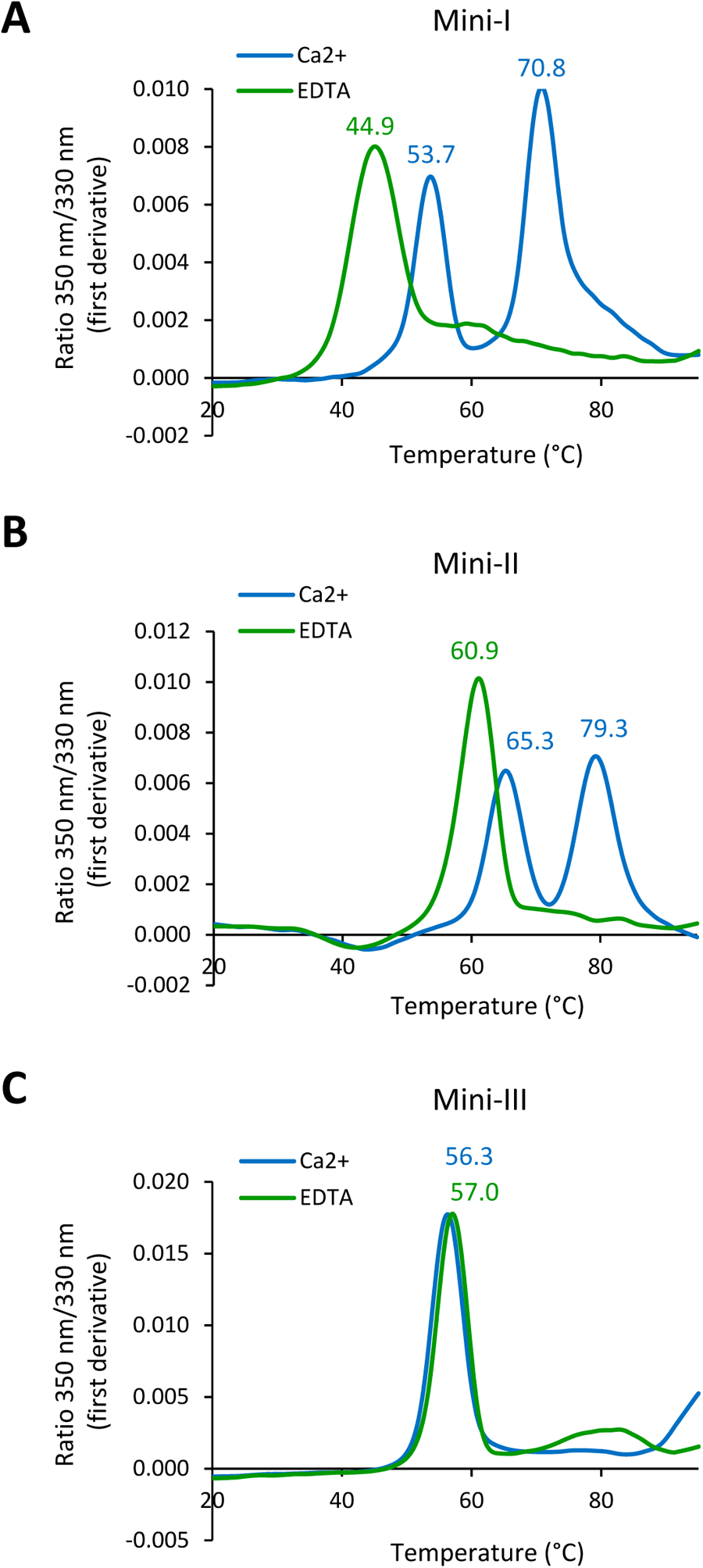
Analysis of the thermal stability of mini-procollagens in the presence and absence of calcium by NanoDSF. Fluorescence emitted at 350 nm and 330 nm was recorded during the thermal denaturation of the mini-procollagens in HEPES buffer containing 5 mM CaCl_2_ (Ca^2+^) or 20 mM EDTA. The first derivatives of the 350 nm / 330 nm ratio are shown for Mini-I **(A)**, Mini-II **(B)** and Mini-II **(C)**. Curves are means of two or three experiments performed in duplicate. The corresponding melting temperatures (Tm) are indicated above the curves.

Treatment with EDTA induced a major change of the denaturation curves of Mini -I and -II, with a decrease of Tm1 and the disappearance of Tm2 indicating a global decreased stability of these proteins in the absence of calcium. The effect was more pronounced for Mini-I with a 9° C shift of Tm1 (5°C for Mini-II). Mini-III was not affected, with no change of Tm1. Thus, the thermal stability of Mini-I and II was clearly linked to the presence of calcium, whereas the latter seemed to have much less impact on Mini-III.

### Cleavage of mini-procollagens by BMP-1 and interaction with PCPE-1

The next step was to validate the ability of BMP-1 to efficiently cleave Mini-I and Mini-II and of PCPE-1 to enhance their cleavage. To do this, we incubated the mini-procollagens in a 1:25 enzyme:substrate molar ratio for 15 min at 37°C and analyzed the cleavage products by SDS-PAGE in both non-reducing and reducing conditions (Fig. 3A). All recombinant mini-procollagens could be cleaved by BMP-1 and this cleavage was stimulated by PCPE-1 (here in a 1:1 molar ratio to mini-procollagen), as expected. Based on the amount of C-propeptide in reducing conditions (lower panel in Fig. 3A), it seemed that all mini-procollagens showed similar extents of cleavage, both with and without PCPE-1. Interestingly, after BMP-1 cleavage, the two bands corresponding to the cleavage products of the α1 and α2 chains of Mini-I were clearly resolved by SDS-PAGE and confirmed the heterotrimeric nature of Mini-I with the correct 2:1 ratio. Also, the cleavage pattern in non-reducing conditions (upper panel in Fig. 3A) indicated that processing occurs through intermediate steps with only one chain or two chains cleaved before the C-propeptide can be fully released.

**Figure 3:**
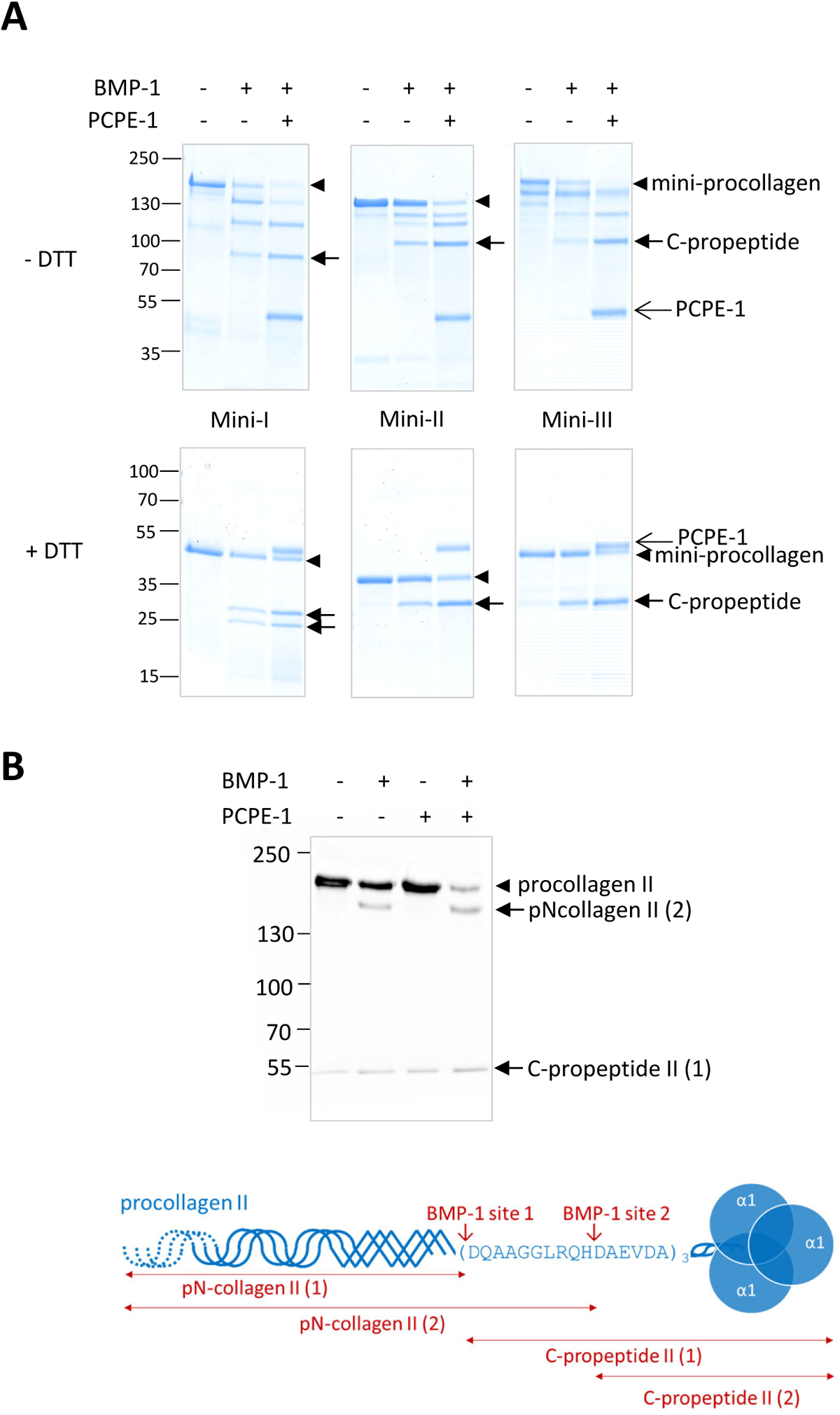
Validation of the use of mini-procollagens to analyze the PCP activity of BMP-1 and its enhancement by PCPE-1. **(A)** Cleavage assays of the three mini-procollagens (450 nM) in the absence or presence of BMP-1 (18 nM) or in the presence of BMP-1 (18 nM) and PCPE-1 (450 nM) for 15 min. Analysis was by SDS-PAGE in non-reducing (upper panel) or reducing (lower panel) conditions (4-20 % acrylamide gradient gel, InstantBlue staining). The gels are representative of three independent experiments. **(B)** Immunodetection of cleavage products derived from native procollagen II in the supernatant of articular chondrocytes incubated (i) alone or (ii) with 10 nM BMP-1 or (iii) with 500 nM PCPE-1 or (iv) with 10 nM BMP-1 + 500 nM PCPE-1 (as indicated above the gel) for 3 h at 37°C (8 % acrylamide gel, reducing conditions). The blot is representative of three independent experiments performed in duplicates with the supernatant of two different donors.

The enhancing activity of PCPE-1 was previously found to depend on its tight interaction with C-propeptides, as demonstrated for native C-propeptide I and for recombinant C-propeptide III (15, 19, 25). Therefore, we also analyzed the binding of Mini-I and -II to immobilized PCPE-1 by Surface Plasmon Resonance (SPR) and compared the results with the binding of Mini-III (Fig. 4A). Interestingly, all mini-procollagens could interact with PCPE-1 but the curve profiles were significantly different. Mini-I and -III had similar association rates but Mini-I formed a more stable complex with PCPE-1, as demonstrated by its lower dissociation rate. Mini-II had a completely different behavior with both association and dissociation being faster than for the other two mini-procollagens, indicating efficient but transient binding. These findings were confirmed by successive injections of serial dilutions of the various mini-procollagens (Fig. 4B-D) and curve fitting with the heterogenous ligand model which was previously found to give the best results with Mini-III (Table 1). This model yielded two dissociation constants (K_D1_ and K_D2_ with K_D2_ contributing for the most important part, between 73 and 85 %, of the maximum signal or R_max_) which were all below 15 nM indicating that all mini-procollagens were strong ligands of PCPE-1. In agreement with the different profiles observed in Fig. 4A, Mini-II displayed the fastest association rates (k_a1_ and k_a2_) and Mini-I the slowest dissociation rates (k_d1_ and k_d2_) (Table 1).

**Figure 4:**
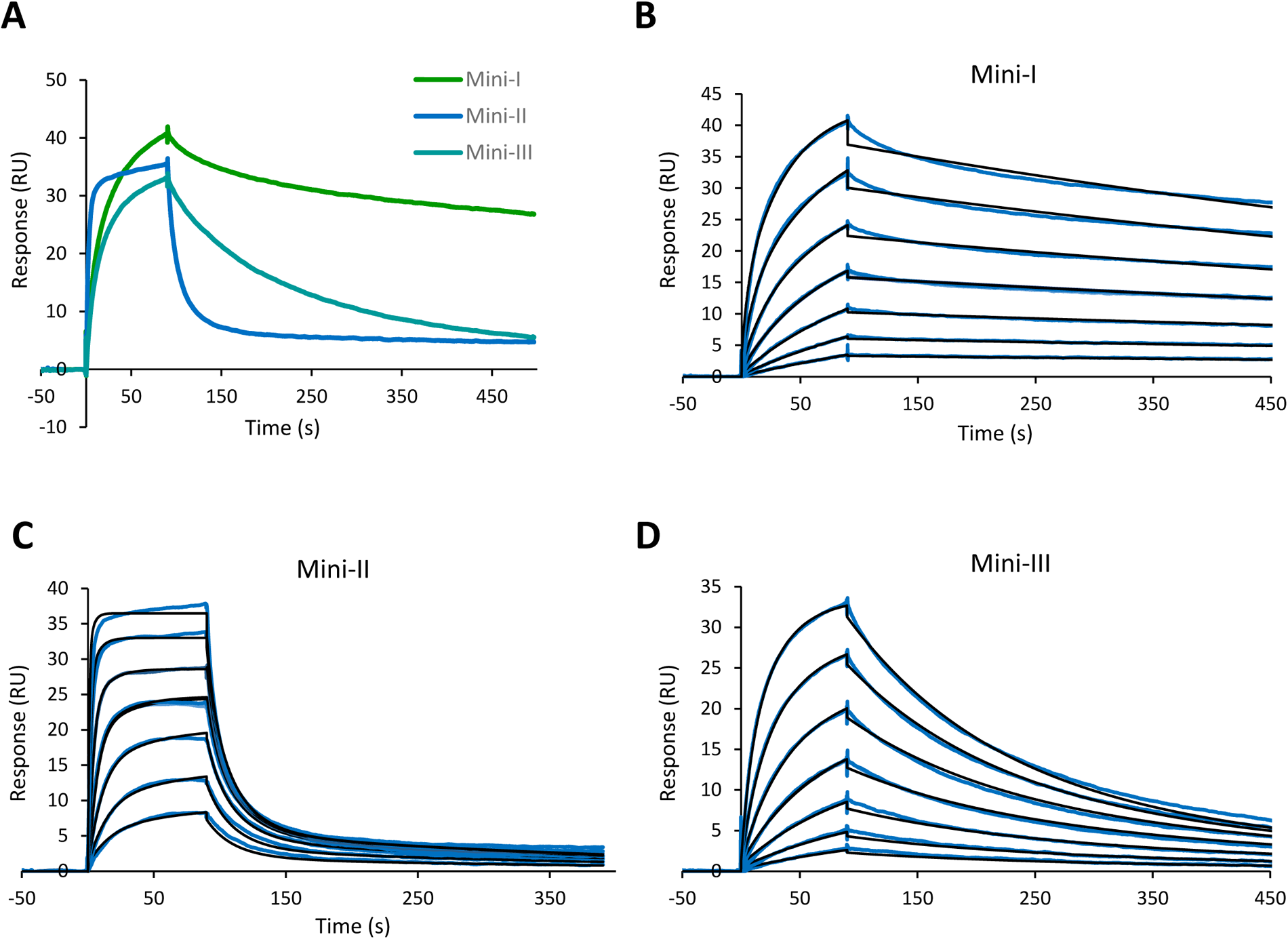
Interaction of mini-procollagens with PCPE-1 analyzed by SPR. **(A)** Injection of 50 nM of each mini-procollagen over immobilized PCPE-1 (70 RU). **(B, C, D)** Increasing concentrations of Mini-I (A), Mini-II (B) and Mini-III (C) were injected (0.78-50 nM, prepared as serial two-fold dilutions, with one duplicate injection of 6.25 nM mini-procollagen). The best fit of the experimental curves obtained with the heterogenous ligand model is also shown.

**Table 1:**
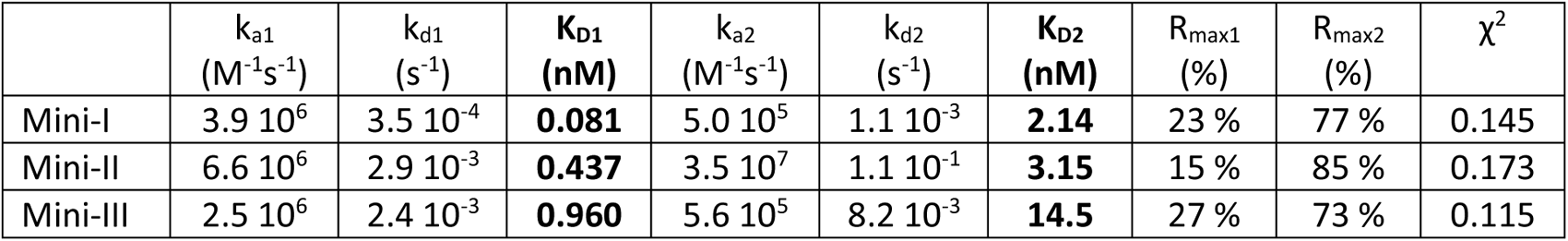
Dissociation constants for the interaction of immobilized PCPE-1 with mini-procollagens determined by SPR using the heterogeneous ligand model. Experimental curves and their best fit are shown in Figure 4.

All these results validated the use of mini-procollagens as model substrates to analyze BTP procollagen C-proteinase activity and to gain mechanistic information on the regulation of the procollagen maturation complex by PCPE-1 or other regulatory proteins.

### Cleavage site identification in mini-procollagens

We then decided to confirm the previously described cleavage sites for BMP-1 in procollagens which were determined a long time ago (26–28) with potentially less sensitive techniques. We used a mass-spectrometry-based approach called ATOMS (Amino-Terminal Oriented Mass spectrometry of Substrates) which relies on the specific isotopic labeling of N-termini, either naturally present in proteins or generated by proteolysis (29, 30). Mini-procollagens were incubated alone, with BMP-1 or with BMP-1 and PCPE-1 and the last two conditions were compared to the control condition without protease. Cleavage sites were revealed by N-terminally-labeled peptides with isotopic ratios superior to 2. Interestingly, using this approach, we confirmed all the previously described cleavage sites (marked with an asterisk in Table 1) and they all had the canonical aspartate in P1’ position. However, we also found one additional cleavage in the α1 chain of Mini-I and of Mini-II, suggesting that previous information was not complete. These new cleavages occurred 7 or 10 residues further down previously described sites respectively and also had aspartate in P1’ position. Importantly also, the same cleavage sites were identified in the absence and presence of PCPE-1, indicating that PCPE-1 does not change the cleavage specificity of BMP-1.

The ATOMS mass spectrometry experiment was performed with hydroxy-proline as variable modification to allow the best coverage of the triple helix in mini-procollagens. A sequence coverage of 100 % was actually obtained for the triple helices of Mini-I and Mini-II and of 70 % for Mini-III. No BMP-1 cleavage site was found in this part of the recombinant proteins despite the presence of several aspartate residues, suggesting that they were properly folded. The latter conclusion was also supported by the finding that most prolines in recombinant mini-procollagens could be hydroxylated with at least one peptide occurrence showing their hydroxylation (data not shown).

Finally, the presence of a second BMP-1 cleavage site in procollagen II could be confirmed by the analysis of the conditioned medium of articular chondrocytes after incubation with BMP-1 or BMP-1 + PCPE-1. Using immunoblotting with an antibody directed against the DQAAGGLRQHDAEVDA peptide (31) which starts like the C-terminal product defining the proximal (or canonical) cleavage, we detected a band matching the expected size of C-propeptide II but also a high-molecular weight product which likely corresponds to a pN-collagen form still containing the DQAAGGLRQH sequence after cleavage at the second site (Fig. 3B). This result strongly suggests that cleavage by BMP-1 at the two adjacent sites can also occur in native full-length procollagen II.

## Discussion

In the present study, we describe the successful production of heterotrimeric mini-procollagen I and of homotrimeric mini-procollagen II. Our system proved to be very versatile and allowed the production of both homotrimers and heterotrimers with the correct stoechiometry. Interestingly also, mini-procollagens could be obtained in the absence of N-glycosylations, with different lengths of the triple helical region and different tags (6His, Twinstrep, c-myc) without disrupting the overall structure of the molecules. The yield was better for Mini-II than for Mini-I, a result that can be explained by the fact that the α1(I) chain forms homotrimers in addition to the heterotrimers. It could be interesting to exploit this property to simultaneously purify the homotrimers but the addition of another tag on the α1(I) chain would then be needed. Notably, the overall yields obtained with transient transfections of HEK293-F cells were substantially higher than the yield previously obtained with Mini-III through stable transfection of HEK293-EBNA cells. A recent test to produce mini-procollagen III using the same type of construct and the same production protocol as those described here for Mini-II showed that the yield could also be increased by a factor of at least ten (data not shown), suggesting that the new system is significantly better than the previous one, regardless of mini-procollagen sequence.

An important feature of our production system is that it allows proline hydroxylation in the collagen triple helices, an important determinant of their stability at 37 °C. Analysis of the thermal denaturation of the mini-procollagens also confirmed their good overall thermal stability but suggested different dependencies upon calcium binding, a result that will require further investigation in the future.

The availability of a complete toolkit of mini-procollagens derived from the three major fibrillar collagens now offers a very large range of perspectives to make functional and structural studies. Here, we have shown the potential of the mini-procollagens for cleavage site identification with BMP-1 and this could be further extended to more recently-evidenced PCPs such as meprins (16), ADAMTS-2 and ADAMTS-14 (17). Mechanistic studies are now also made possible, especially through site-directed mutagenesis, to characterize the molecular determinants of procollagen interactions with proteolytic enzymes or various potential partners. Here, we have characterized the interaction of the three mini-procollagens with PCPE-1, one of their best-studied partner involved in the control of their proteolytic maturation by BTPs (32), meprins (33) but not ADAMTSs (17). Intriguingly, even if the overall affinity of the three mini-procollagens for PCPE-1 was quite strong (K_D_ in the low nanomolar range), they displayed very different association and dissociation kinetics that would be worth exploring in more detail as they probably reveal distinct structural features. Finally, based on the suitability of the mini-procollagens for cleavage and SPR assays, it should also be possible to establish medium-throughput screening assays to test small-molecule compounds or antibody-related inhibitors susceptible to modulate their activities.

One interesting application of the mini-procollagens described here is the easy determination of cleavage sites for various proteases. In this context, we were able to confirm previously reported cleavage sites for BMP-1 but also to reveal one additional cleavage in the α1 chain of procollagen I and in procollagen II, occurring some residues downstream of the canonical sites. It cannot be excluded that this is an artefact of the recombinant system leading to partially unfolded proteins but, at least in the case of procollagen II, we have good evidence that this second cleavage also occurs in the native molecule. First, we observed that an antibody directed against the beginning of the canonical C-propeptide II could also detect a pN-collagen-like product, suggesting the conservation in some of the BMP-1-generated pN-collagen molecules of the sequence recognized by the antibody (Fig. 3B). Then, it has been described by Van der Rest and colleagues that C-propeptide II (also known as chondrocalcin) is present in two forms in the bovine cartilage extracellular matrix (34). The sequences of the two forms exactly match the two cleaved forms of C-propeptide II identified here in the presence of BMP-1, except at species-specific positions. Interestingly, in fœtal epiphyseal cartilage, the shortest form (called C-propeptide II (2) in Fig. 3B) is twice more abundant than the longest form (C-propeptide II (1)) (34), suggesting that this cleavage site is also fully relevant *in vivo*. Together, the results also indicate that the two cleavage events occur independently rather than sequentially as it does not seem that any of the cleavages is required for the second cleavage to happen.

In conclusion, the newly developed mini-procollagens already appear as valuable tools to analyze the maturation process of fibrillar collagens in mechanistic and, potentially also, structural studies. They could also have great potential for the development of screening assays in the context of anti-fibrotic drug development (35).

## Materials and methods

### Mini-procollagen production

The DNA sequences encoding the two chains of mini-procollagen I with supplementary GPP repeats and tags but without signal peptide, as described in Fig. S1, were synthesized by Genewiz using native sequences with mutations of N-glycosylation sites. Integration into the pHLsec vector was also done by Genewiz using the AgeI and XhoI cloning sites, in frame with the signal peptide already included in the vector (24).

The mini-procollagen II construct, as described in Fig. S1, was obtained by PCR amplification of the procollagen II sequence found in human chondrocyte cDNAs using GGACCACCAGGGCCTCCTGGTCCCCGTGGA and ATCGCGATACTAGTCTGGGTTCAGGTTTTTACAAGAAGCAACGGATTGTG as forward and reverse primers respectively. This DNA fragment together with another sequence encoding the N-terminal 6His tag and an additional (GPP)_4_ sequence, obtained by hybridizing the following primers: GTTGCGTAGCTGAAACCGGTCATCATCACCACCATCACGGACCACCAGGACCACCAGGACCACCAGGACCACCAGGGCCT (forward) and AGGCCCTGGTGGTCCTGGTGGTCCTGGTGGTCCTGGTGGTCCGTGATGGTGGTGATGATGACCGGTTTCAGCTACGCAAC (reverse), were then cloned into pHLsec, digested with AgeI and XhoI, using the Gibson Assembly cloning kit (NEB) according to manufacturer’s protocols.

Insert sequences were verified by Sanger sequencing (Eurofins) before plasmid amplification in *E. coli* XL1-Blue bacteria. Plasmids were then purified using the NucleoBond PC 10000 EF Giga kit for endotoxin-free plasmid DNA from Macherey-Nagel. In parallel, HEK293-F cells were grown in FreeStyle 293 expression medium (Gibco) (20) using sterile flasks (Corning or BD Biosciences) placed on an orbital shaker platform rotating at 125 rpm at 37 °C with 8 % CO_2_ (Eppendorf). On the day of transfection, cells were centrifuged and resuspended at a cell density of 1.10^6^ cells/mL in FreeStyle 293 expression medium. For transfection, the ratio of total DNA (in a 2:1 mass ratio for the α1 and α2 chains of Mini-I) to cells was 1 µg DNA for 10^6^ cells and the ratio of total DNA to PEI Max transfection agent (Polyethylenimine HCL MAX 40 K linear, Polysciences) for 1 µg DNA for 3 µg transfection agent. Plasmid DNA and PEI Max transfection agent were first diluted separately in 1/20 of the total culture volume of Opti-MEM medium (Gibco), filtered on 0.2 µm filters (Millipore) for the transfection agent and kept at room temperature for 5 min before mixing. The mixture was further incubated for 15 min before addition to the cells. Ascorbic acid (50 µg/mL final concentration) was also added to the culture. Finally, 1/5 of the culture volume of fresh medium was added to the cells 24 hours after transfection.

After 4 days, the suspension was centrifuged at 1,000 g for 10 min, the cells were discarded and *N*-ethyl-maleimide (Merck, 2 mM) and Pefabloc (Roth, 0.25 mM) were added to the culture medium before centrifugation for 10 min at 10,000 g, 4° C. The pH of the supernatant was adjusted to 8. In the case of Mini-I, a biotin-blocking solution (IBA lifesciences) was added and incubated with the medium for 15 min at 4 °C under gentle stirring. The cell supernatants were frozen at -80 °C until purification.

Mini-III was produced in HEK293-EBNA cells, as previously described (14).

### Mini-procollagen purification

The purification of Mini-I was achieved in two steps using the Twinstrep tag on the α2 chain and size exclusion chromatography. After equilibration of the Strep-tactin XT Superflow resin (IBA lifesciences, 10 mL per liter of cell supernatant) in 0.1 M Tris-HCl pH 8, 0.3 M NaCl, the cell supernatant was loaded at 0.5 mL/min, 4 °C. The column was washed with 5 volumes of equilibration buffer and eluted with 6 volumes of the same buffer supplemented with 50 mM D-biotin (IBA lifesciences). The fractions containing Mini-I were pooled and concentrated by ultrafiltration (Vivaspin 20, 30 kDa, Sartorius) before injection in a HiLoad 10/300 Superdex S200 Increase gel filtration column (Cytiva) equilibrated in 20 mM HEPES pH 7.4, 0.15 M NaCl, 5 mM CaCl_2_. Useful fractions were concentrated around 1 mg/mL and stored at -80 °C until further use.

The purification of Mini-II was also achieved in two steps but the first step was with Ni Sepharose Excel (Cytiva) chromatography. Cell supernatant was loaded on the resin (3 mL per liter of supernatant), equilibrated in 20 mM HEPES pH 8, 0.3 M NaCl (buffer A), at 1 mL/min. Then, the resin was washed successively with buffer A and buffer A containing 25 mM imidazole and the protein was eluted with a gradient of imidazole from 25 to 500 mM. Fractions containing Mini-II were pooled and submitted to SEC in the same conditions as above except that the running buffer was 20 mM HEPES pH 7.4, 0.3 M NaCl, 2.5 mM CaCl_2_.

The purification of Mini-III was performed as previously described (14) with some modifications. Affiblue resin (Bio-Rad) was used instead of Heparin sepharose and eluted with PBS, 2 M NaCl. The fractions containing Mini-III were then loaded through an anti-PCPE-Sepharose (prepared by cross-linking the purified polyclonal antibody directed against PCPE-1 (14) to CNBr-activated Sepharose 4B from Cytiva) to remove endogenous PCPE-1 and further immunopurified on an anti-c-Myc column, as described (14). A final step on HiLoad 10/300 Superdex S200 Increase gel filtration column (Cytiva) was also added with 20 mM HEPES pH 7.4, 0.15 M NaCl, 5 mM CaCl_2_ as running buffer.

### Other proteins

Human BMP-1 and PCPE-1 (with or without His tag at the C-terminus) were obtained as described before (14, 36, 37).

### Deglycosylation with PNGase F

Removal of N-linked oligosaccharides in mini-procollagens was performed with PNGase F (New England BioLabs, glycerol-free). Ten micrograms of mini-procollagens were first denatured at 100 °C for 10 min in Glycoprotein Denaturing Buffer provided with the enzyme. The deglycosylation reaction was then carried out at 37 °C for 4 h in GlycoBuffer 2 with 1 % NP-40 and 70 U PNGase F and analyzed by SDS-PAGE in reducing conditions.

### Western blotting

After SDS-PAGE, proteins were transferred to polyvinylidene difluoride (PVDF) membranes by liquid transfer in 10 mM *N*-cyclohexyl-3-aminopropanesulfonic acid (pH 11) with 10 % ethanol (2 h, 280 mA) and blocked for 2 hours in 10 % (w/v) skim milk in PBS. Primary antibodies against PCPE-1 (rabbit polyclonal) and procollagen II (mouse monoclonal) were home-made antibodies, prepared as described in (14) and (31) respectively. They were diluted in 5 % skim milk, 0.05 % Tween 20 in PBS and applied on membranes overnight at 4°C. After three washes in 0.05 % Tween 20 in PBS, horseradish peroxidase (HRP)-coupled secondary antibodies (Cell Signaling Technology, 1:10,000) were added for 1 h at room temperature. Signal was detected after extensive washing using the enhanced chemiluminescence technique with the ECL Select kit (Cytiva) and the FX Fusion camera (Vilber Lourmat).

### Cleavage assays

Mini-procollagens were incubated with BMP-1, in the presence of absence of PCPE-1 (with an 8His-tag at the C-terminus (36)), at 37°C under for the time indicated in figure and table legends. The buffer was 50 mM HEPES pH 7.4, 150 mM NaCl, 5 mM CaCl2, 0.02 % octyl β-D-glucopyranoside. The reactions were started by the addition of the protease and stopped with 25 mM EDTA and/or SDS-PAGE sample buffer. They were analyzed by SDS-PAGE on Criterion gradient gels (Bio-Rad) with InstantBlue Coomassie Protein Stain (Abcam) or by ATOMS.

### NanoDSF

Thermal denaturation was analyzed by Nano-DSF using a Prometheus NT.48 instrument (Nanotemper). Mini-procollagens were diluted to a final concentration of 150 µg/mL in 20 mM HEPES pH 7.4, 150 mM NaCl, 5 mM CaCl_2_ (Ca^2+^ condition) or 20 mM HEPES pH 7.4, 150 mM NaCl, 20 mM EDTA (EDTA condition). Prior to dilution, the storage buffer of Mini-II was exchanged on a Zeba-spin column (Thermo Scientific) for 20 mM HEPES pH 7.4, 150 mM NaCl, 5 mM CaCl_2_. Ten microliters of proteins were loaded into a capillary (PR-C002, Nanotemper) and thermal denaturation was recorded using a 1 °C/min ramp from 20 to 95 °C, with fluorescence detection at 330 nm and 350 nm (excitation at 280 nm). Data were recorded using the PR.ThermoControl v2.3.1 software and analyzed with the PR.StabilityAnalysis v1.1 software (Nanotemper). Melting temperatures were determined as the inflection points of the curves representing the ratio of fluorescence intensity at the two wavelengths (350nm/330nm) as a function of temperature.

### Surface Plasmon Resonance

Surface plasmon resonance experiments were run at 25 °C with a Biacore T200 equipment (Cytiva) equipped with the Biacore T200 control software (v3.2.1). Human PCPE-1 (no tag (14)) was diluted in 10 mM HEPES pH 7.4 and covalently immobilized on Series S sensor chips CM5 (Cytiva) using the reagents included in the Amine coupling kit (Cytiva). SPR signals were recorded simultaneously on a control channel where a mock immobilization procedure was performed (no protein ligand) and double referencing was applied to all injections. Mini-procollagens were injected at 30 μl/min after dilution in running buffer (10 mM HEPES pH 7.4, 0.15 M NaCl, 5 mM CaCl2 and 0.05 % P20) for 90 s. Regeneration was achieved with 2 M guanidinium chloride. Best fits of the data were computed with the Biacore T200 evaluation software (v3.2.1).

### Chondrocyte cultures

Human articular chondrocytes were isolated from macroscopically healthy zones of osteoarthritic knee joints obtained from two donors undergoing total knee replacement. The study was performed in full compliance with local ethics guidelines, national and European Union legislation regarding human sample collection, manipulation and personal data protection. The study protocol was approved by the ethics committee of COnservation D’Eléments du COrps Humain (CODECOH: DC-2014-2325) for preservation of and research with human samples. The cartilage samples were collected after written informed consent of donors. Briefly, small slices of cartilage were digested in a culture medium consisting of Dulbecco’s modified Eagle medium/Ham’s F12 (Gibco) with 0.06 % bacterial collagenase A (Roche Applied Science) overnight. The cells were then seeded at a density of 1.5 x 10^4^ cells/cm^2^ on culture dishes with culture medium supplemented with 10 % fetal bovine serum (FBS, Gibco), 100 μg/mL streptomycin and 100 U/mL penicillin (Thermo Scientific). Forty-eight hours after seeding, the medium was changed for serum-free medium and supplemented with 50 µg/mL sodium ascorbate. The cell supernatant was collected after 2 days and used for cleavage assays on procollagen II.

### ATOMS

Five micrograms of cleaved and uncleaved mini-procollagen samples, prepared as above, were processed for TMT-ATOMS, as previously described (29, 30) with minor modifications. Briefly, samples were denatured in 2.5 M guanidinium chloride and 1.8 M HEPES pH 8.0 at 65 °C for 15 min, reduced with 10 mM Tris (2-carboxyethyl) phosphine (TCEP) at 65 °C for 45 min, and alkylated with 25 mM iodoacetamide at room temperature in the dark for 30 min. After TMT labeling in DMSO (TMT label:protein mass ratio = 26), the three mini-procollagen samples (alone, with BMP-1, with BMP-1 + PCPE-1) were mixed and concentrated using the SP3 (single-pot, solid-phase-enhanced sample-preparation) protocol (38). Proteins and SP3 beads (GE healthcare) were mixed in a 1:10 weight ratio in 80 % ethanol for 5 minutes at RT. Then, supernatant was discarded and beads were washed three times on a magnetic rack with 80 % ethanol before being resuspended in 100 µL of 180 mM HEPES pH 8. The supernatant was digested overnight at 37 °C with sequencing-grade trypsin or chymotrypsin (Promega; 1:100 protease:protein weight ratio). After desalting on a C18 column (Pierce) and drying under vacuum, the digested samples were dissolved in 0.1 % formic acid and analyzed by LC-MS/MS using a Q-Exactive HF mass spectrometer (Thermo Scientific), operated with the Xcalibur software (version 4.0) and equipped with a RSLC Ultimate 3000 nanoLC system (Thermo Scientific). Data files were analyzed with Proteome Discover 2.4 using the SEQUEST HT search engine against a homemade database including the mini-procollagens, BMP-1 and PCPE-1 sequences. Precursor mass tolerance was set at 10 ppm, fragment mass tolerance at 0.02 Da and up to 3 missed cleavages were allowed. Semi-tryptic digestion was selected. Oxidation (M, P), acetylation (Protein N-terminus), and TMT6Plex (N-term, K) were set as variable modifications and carbamidomethylation (C) as fixed modification. Peptides and proteins were filtered using the XCorr confidence threshold calculation based on the ion precursor charges and the MS/MS spectra. Quantification was performed with the Reporter Ions Quantifier node. The peak integration was set to the Most Confidence Centroid with 20 ppm Integration Mass Tolerance on the reporter ions. Protease/control ratios were calculated as the means of all the ratios obtained for N-terminally-labeled peptides defining cleavage sites (regardless of C-terminal length or modifications) or mature N-termini (after signal peptide removal). Ratios showing a fold change of at least two were conserved as defining cleavage sites in Table 2.

**Table 2:** Cleavage site identification by ATOMS. Mini-procollagen substrates (950 nM) were incubated either with 46 nM BMP-1 or with 24 nM BMP-1 and 950 nM PCPE-1 for 1 h and, after TMT-labeling, their N-terminal peptides were compared to the control condition (without BMP-1 and PCPE-1). ATOMS ratios were calculated as the means of all the ratios obtained for peptides defining cleavage sites (↓) or mature N-termini (after signal peptide removal) of mini-procollagens. Cleavage site positions are indicated as in reference human proteins identified by their Uniprot ID and previously described cleavage sites are marked by an asterisk (*). The working enzyme used to digest proteins could be trypsin or chymotrypsin.

| Reference protein | Working enzyme | Mature N-terminus or cleavage site | Protease/control ratio (number of peptide identifications) |  |
| --- | --- | --- | --- | --- |
|  |  |  | BMP-1 | BMP-1 + PCPE-1 |
| Collagen I (α1)<br>P02452 | chymotrypsin / trypsin | Mature N-terminus in Mini-I (α1) | 0.8 (11) | 0.5 (11) |
|  | chymotrypsin / trypsin | YYRA↓<sup>1219</sup>DDAN* | 4.4 (8) | 6.9 (8) |
|  | chymotrypsin / trypsin | NVVR↓<sup>1226</sup>DRDL | 3.1 (7) | 10.4 (7) |
| Collagen I (α2)<br>P08123 | chymotrypsin / trypsin | Mature N-terminus in Mini-I (α2) | 0.6 (3) | 0.2 (3) |
|  | chymotrypsin | FYRA↓<sup>1120</sup>DQPR* | 6.7 (7) | 5.1 (7) |
| Collagen II (α1)<br>P02458 | trypsin | Mature N-terminus in Mini-II (α1) | 0.8 (3) | 1.7 (3) |
|  | trypsin | YMRA↓<sup>1242</sup>DQAA* | 7.9 (33) | 5.4 (33) |
|  | chymotrypsin / trypsin | LRQH↓<sup>1252</sup>DAEV | 7.7 (12) | 11.6 (11) |
| Collagen III (α1)<br>P02461 | chymotrypsin | Mature N-terminus in Mini-III (α1) | 1.4 (1) | 0.5 (1) |
|  | chymotrypsin / trypsin | PYYG↓<sup>1222</sup>DEPM* | 4.5 (23) | 4.9 (25) |

## Acknowledgements

We thank Frédéric Delolme and Adeline Page from the Protein Science Facility of UMS 3444 (SFR Biosciences, Lyon) for performing mass spectrometry analyses. We also thank Frédéric Mallein-Gerin and Alexandre Dufour for preparing chondrocyte medium.

---

**Supplementary Figure S1:**
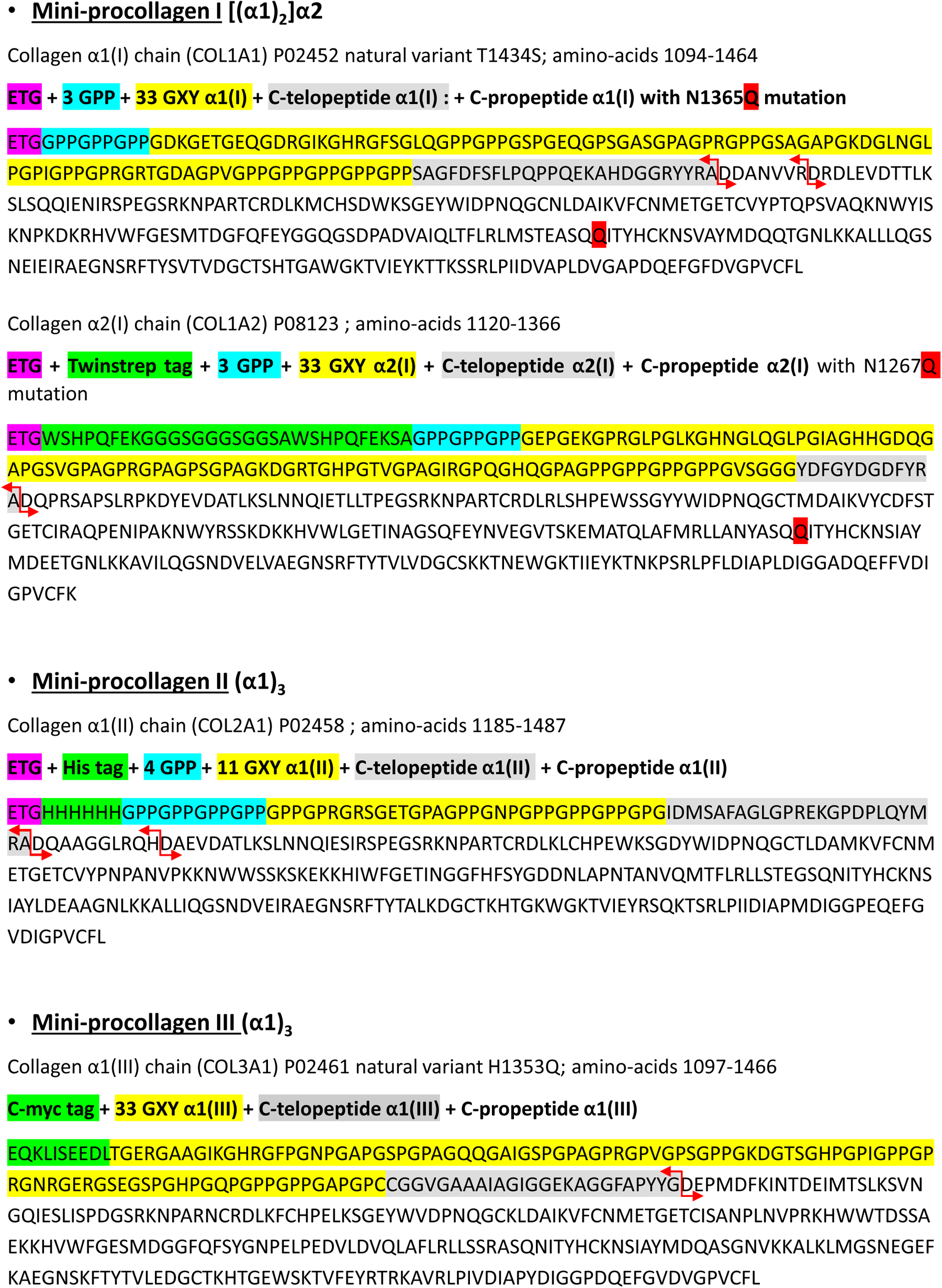
Mature protein sequences for the three mini-procollagens. The different parts of the sequences are shown in different colors (color code as indicated above each sequence); the cleavage sites for BMP-1, identified in this study, are indicated by red arrows.

